# Individual-based modeling of genome evolution in haplodiploid organisms

**DOI:** 10.1101/2021.10.25.465450

**Authors:** Rodrigo Pracana, Richard Burns, Robert L. Hammond, Benjamin C. Haller, Yannick Wurm

**Author notes:** Joint first authors (Rodrigo Pracana, Richard Burns).

## Abstract

Ants, bees, wasps, bark beetles, and other species have haploid males and diploid females. Although such haplodiploid species play key ecological roles and are threatened by environmental changes, no general framework exists for simulating their genetic evolution. Here, we use the SLiM simulation environment to build a novel model for individual-based forward simulation of genetic evolution in haplodiploid populations. We compare the fates of adaptive and deleterious mutations and find that selection is more effective in haplodiploid species than in diploid species. Our open-source model will help understand the evolution of sociality and how ecologically important species may adapt to changing environments.

## Main text

Approximately 15% of all animal species, including ants, bees, wasps, thrips, and bark beetles, have a haplodiploid sex-determination system: haploid, unfertilized eggs develop into males, while diploid eggs develop into females (Pamilo and Crozier 1981). These haplodiploid species display a huge diversity of morphologies and behaviors. For example, solitary and social bees occupy essential ecological and agricultural roles as pollinators (Potts et al, 2016), and bees and ants are charismatic models for studying social evolution (Hölldobler and Wilson, 1990).

Haplodiploid populations evolve differently from diploid populations (Hedrick and Parker, 1997), yet no flexible, general genome simulation framework incorporates haplodiploidy. Simulations are crucial for understanding how demographic and genomic processes affect haplodiploid evolution, enabling the interpretation of genome-wide analyses of haplodiploid species, testing competing hypotheses regarding their abilities to respond to human-induced environmental changes, and exploring genomic models of social evolution (Bank et al. 2014; Favreau et al 2018; Lopez-Osorio and Wurm 2020; Colgan et al. 2021).

Here, we present a model for individual-based forward evolution of haplodiploid genomes. Our model is built upon SLiM, a software framework for genomic simulation of populations of individuals (version 3.3; Haller and Messer, 2019). To work around SLiM’s assumption that all individuals are diploid, we restrict males to inherit one random chromosome from the recombined genome of a female and give them a NULL second chromosome. We assign a relative fitness of 1 + *s* to male carriers of a mutation with selection coefficient *s*. The Supplementary Text includes more details. Our model can be extended to consider variation in recombination rates, selection, and complex demographic structures.

We used our model to compare the fates of mutations between haplodiploid populations and diploid populations. We first subjected both types of population to neutral mutations (*s* = 0). As expected from evolution by drift, with a mutation rate of 10^−8^ on a genome of 10^6^ loci, 0.01 mutations were fixed per generation in both population types (fig. 1A). Because recessive mutations are fully exposed to selection in haploid males but masked in heterozygote males of diploid populations, we subsequently tested whether selection on recessive mutations is more effective in haplodiploid populations (Avery 1984). This was indeed the case. Advantageous recessive mutations (*s* > 0) fixed at a higher rate in haplodiploid populations, with stronger effects in simulations with larger selection coefficients (fig. 1B). Similarly, weakly deleterious recessive mutations (*s* = −0.0005 and *s* = −0.001) fixed at a lower rate in haplodiploid populations (fig. 1C). Very few more strongly deleterious mutations (*s* = −0.003 and *s* = −0.01) became fixed in either type of population, likely because selection against such mutations can overpower drift.

**FIG. 1.**
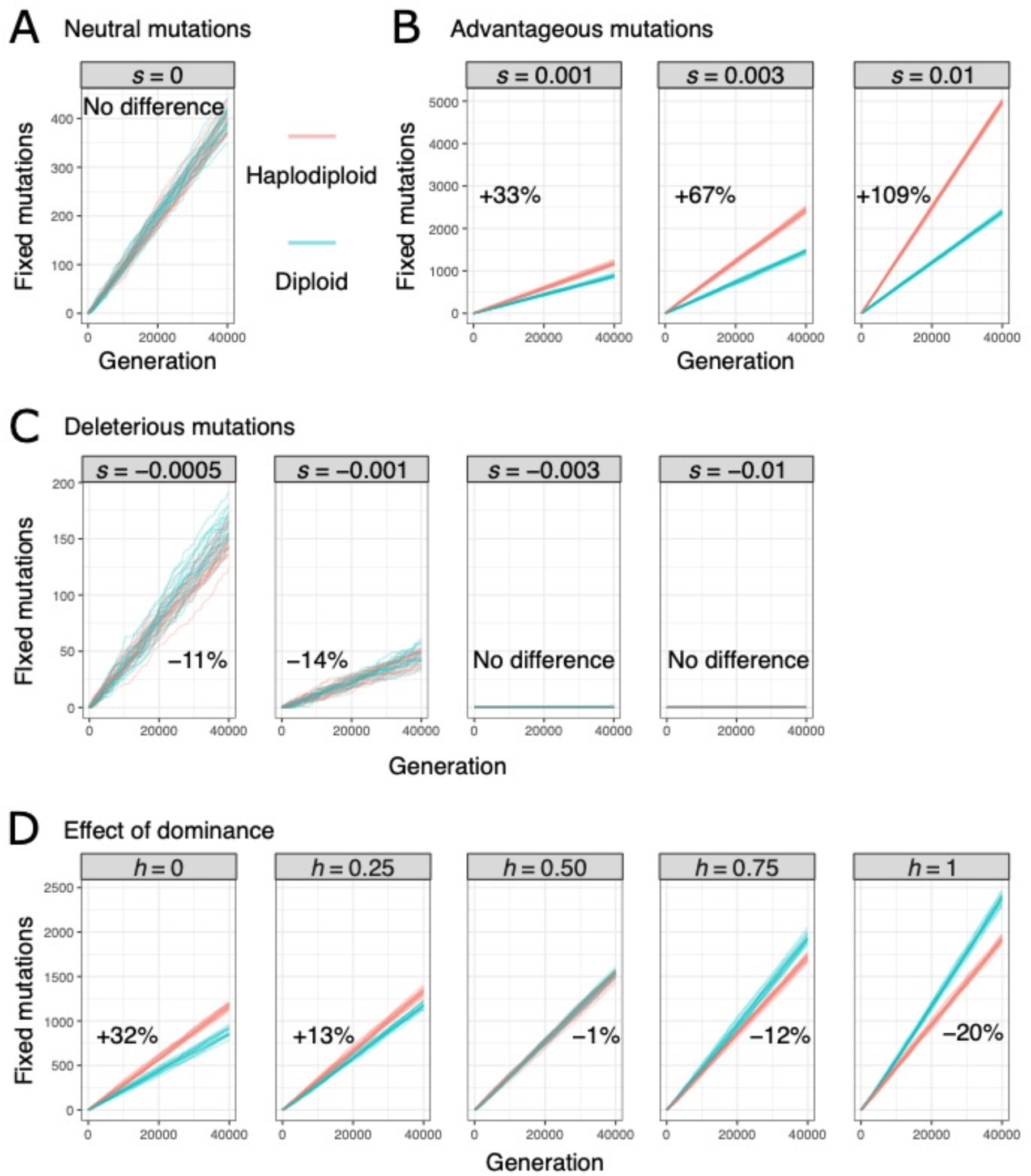
The effect of haplodiploidy on the fixation rate of (A) neutral mutations, (B) advantageous mutations, (C) deleterious mutations, and (D) advantageous mutations with different levels of dominance. Each line represents one simulation run (only 20 shown for each treatment). On each plot, we also show the average difference in the number of fixed mutations between haplodiploid and diploid simulations after 40,000 generations and a burn-in period of 10,000 generations, when statistically significant (Wilcoxon rank sum test, *p* < 0.05; Supplementary Table 1). For each treatment, we ran 200 simulations with populations of 1,000 males and 1,000 females, a genome with 10^6^ loci, a mutation rate of 10^−8^, and a recombination rate of 10^−6^. For the simulations in A to C, mutations were fully recessive (dominance coefficient *h* = 0) and had a range of selection coefficients (*s*, as shown); for the simulations in D, mutations had *s* = 0.001 and a range of dominance coefficients (*h*, as shown).

In reality, most mutations are neither fully recessive (dominance coefficient *h* = 0, as considered in the simulations described so far) nor fully dominant (*h* = 1). We therefore compared fixation rates of advantageous mutations between population types across a range of dominance coefficients. For simulations of recessive mutations (*h* < 0.5), selection was more effective in haplodiploid populations. However, when mutations were dominant (*h* > 0.5), this pattern was reversed (fig. 1D). This somewhat counterintuitive reversed pattern occurs because haplodiploid populations have fewer chromosomes than diploid populations of the same size (1.5*N* versus 2*N*), so they have fewer mutations entering the population in each generation (1.5*Nμ* versus 2*Nμ*), and thus, fewer mutations that can ultimately become fixed (Charlesworth et al. 2018). If both population types have identical numbers of chromosomes rather than individuals, the main pattern of higher efficacy of selection in haplodiploid should prevail whenever *h* < 1.

Overall, our results agree with predictions and empirical evidence regarding selection on X-linked alleles (Charlesworth et al. 2018), which are inherited in a pattern similar to haplodiploidy. The ability to model haplodiploid evolution has the potential to fill major gaps in the study of the demographic and selective processes that have shaped the evolution of social behaviors and other ecologically important traits in ants, bees, and other haplodiploid species. We hypothesize that higher efficacy of selection against recessive deleterious mutations in haplodiploid species may have facilitated the evolution of long lifespans of ant and bee queens, and that it may reduce the degeneration of supergene regions of suppressed recombination (Stolle et al. 2019, Martinez-Ruiz et al. 2020). Additionally, haplodiploidy may modulate the effects of antagonistic selection between sexes or among colony members (Vicoso and Charlesworth 2009, Eyer et al. 2019). We hope to see our model used in the exploration of such processes, and for understanding how interactions between selection efficacy, population size and migration can affect the abilities of haplodiploid species to adapt to environmental changes (Potts et al. 2016).

## Supporting information

SLiM model of haplodiploidy

## Supplementary Material

Supplementary data include the SLiM model of haplodiploidy presented here, an explanatory Supplementary Text, and Supplementary Table 1.

## Acknowledgments

We thank Richard A. Nichols for useful advice. This work was supported by BBSRC (<u>BB/T015683/1</u>) and NERC (<u>NE/L00626X/1</u>, <u>NE/P012574/1</u>). All simulations were run utilizing Queen Mary’s Apocrita HPC facility (http://doi.org/10.5281/zenodo.438045).

## Author Contributions

Conceptualization, YW and RB; Methodology, YW, RB, BCH and RP; Investigation, RP, RB and YW; Writing—Original Draft, YW, RB, and RP; Writing—Review and Editing, YW, RB, BCH, RH and RP; Supervision, YW.

## References

Avery P. 1984. The population genetics of haplo-diploids and X-linked genes. Genet Res. 44:321–341.

Bank C, Ewing GB, Ferrer-Admettla A, Foll M, Jensen JD. 2014. Thinking too positiveã Revisiting current methods of population genetic selection inference. Trends Genet. 30:540–546.

Charlesworth B, Campos JL, Jackson BC. 2018. Faster-X evolution: Theory and evidence from Drosophila. Mol Ecol. 27:3753–3771.

Colgan TJ, Arce AN, Gill RJ, Ramos Rodrigues A, Kanteh A, Duncan EJ, Li L, Chittka L, Wurm Y, unpublished data. Genomic signatures of recent adaptation in a wild bumblebee. bioRxiv https://doi.org/10.1101/2021.05.27.445509, last accessed June 02, 2021.

Eyer P-A, Blumenfeld AJ, Vargo EL. 2019 Sexually antagonistic selection promotes genetic divergence between males and females in an ant. Proc Natl Acad Sci USA. 116:24157–24163.

Favreau E, Martínez-Ruiz C, Rodrigues Santiago L, Hammond RL, Wurm Y. 2018. Genes and genomic processes underpinning the social lives of ants. Curr Opin Insect Sci. 25:83–90.

Haller BC, Messer PW. 2019. SLiM 3: Forward genetic simulations beyond the Wright– Fisher model, Mol Biol Evol. 6:632–637.

Hedrick, P, Parker J. 1997. Evolutionary genetics and genetic variation of haplodiploids and X-linked genes. Annu Rev Ecol Evol Syst. 28:55–83.

Hölldobler B, Wilson EO. 1990. The ants. Cambridge (MA): Belknap Press of Harvard University Press.

López-Osorio F, Wurm Y. 2020. Healthy pollinators: Evaluating pesticides with molecular medicine approaches. Trends Ecol Evol. 35:380–383.

Martínez-Ruiz C, Pracana R, Stolle E, Paris CI, Nichols RA, Wurm Y. 2020. Genomic architecture and evolutionary antagonism drive allelic expression bias in the social supergene of red fire ants. eLife 9:e55862.

Pamilo P, Crozier R. 1981. Genic variation in male haploids under deterministic selection. Genetics 98:199–214.

Potts SG, Imperatriz-Fonseca V, Ngo HT, Aizen MA, Biesmeijer JC, Breeze TD, Dicks LV, Garibaldi LA, Hill R, Settele J, Vanbergen AJ (2016). Safeguarding pollinators and their values to human well-being. Nature. 540:220–229.

Stolle E, Pracana R, Howard P, Paris CI, Brown SJ, Castillo-Carrillo C, Rossiter SJ, Wurm Y. 2019. Degenerative expansion of a young supergene. Mol Biol Evol. 36:553–561.

Vicoso B, Charlesworth B. 2009. Effective population size and the faster-X effect: and extended model. Evolution. 63:2413–2426.

