## Supplementary Text for "Individual-based modeling of genome evolution in haplodiploid organisms"

* Joint first authors

### Supplementary Text

We present the haplodiploidy simulation model in the standard ODD (Overview, Design concepts, Details) format for describing individual-based models (Grimm et al 2006).

#### Overview

<u>Purpose</u>

The model is designed to simulate the most basic rules of haplodiploid inheritance, upon which further investigations into the evolutionary genomic dynamics of haplodiploid species can be built. Under haplodiploid inheritance (arrhenotoky), females are diploid, and males are haploid. Females can reproduce asexually to produce haploid male offspring from unfertilized eggs, or sexually with a haploid male to produce diploid female offspring. The model is built around no specific species or empirical genome sequence, which maintains flexibility. SLiM supports a wide range of evolutionary dynamics for its models, such as migration, admixture, selective sweeps, complex mating schemes, and continuous-space interactions. This model provides a starting point for any future haplodiploid-based work.

<u>State variables and scales</u>

The model has two hierarchical levels: individual and population. Individuals are characterized by several state variables: individual number (an identifier associated with every individual during every SLiM simulation), sex, age (although generations are currently non-overlapping; every individual lives to age 1), and ploidy (haploid males, diploid females). The population is composed of one non-spatial subpopulation of size $K$ with no further variables, although population structure or spatiality could easily be imposed.

<u>Process overview and scheduling</u>

SLiM divides events within each generation cycle into discrete phases that utilize ‘callbacks’: blocks of code which control specific aspects of the simulation. Each generation starts with a `reproduction()` callback, which controls the reproductive events that produce the next generation of offspring. In our model’s `reproduction()` callback, individuals reproduce, with a probability proportional to their relative fitness, until $K$ new individuals are produced. The new offspring (aged 0) are added and the old generation (aged 1) is removed in an `early()` callback. A `late()` callback detects, logs, and removes mutations that have become fixed in the populations as they no longer contribute to relative fitness. Finally, every 100 generations a `late()` callback outputs the generation number and the number of mutations fixed in the simulation thus far.

#### Design concepts

The effect of fitness is modelled explicitly in the `reproduction()` callback, where an individual’s likelihood to be sampled (with replacement) as a parent for the next generation is proportional to its relative fitness. The fitness of an individual is calculated based upon the mutations present in the individual. Females can reproduce sexually or asexually to respectively produce female and male offspring, while males can only reproduce with females to produce female offspring. Outside of reproduction there is no interaction between the individuals in the model. Mutation and recombination rates (per base position, per generation) are taken from constants set in the `initialize()` callback at the beginning of simulation. Stochasticity in the model arises from these mutation and recombination rates, along with the semi-random selection of parents. To validate the model, the rate of mutation fixation for different mutation selection coefficients was compared to theoretical expectations.

### Details

Full code for the model is available as a Supplementary File. Here, we outline the concepts used, in some cases illustrated by code snippets.

Initialization

SLiM models begin with an `initialize()` callback which defines constants and parameterizes the model. In this model's `initialize()` callback the model type is selected to be non-Wright–Fisher so that we can override default reproduction methods. Sex is configured to be modelled explicitly, as otherwise individuals would be hermaphroditic.

We set several parameters that could vary across simulations. A constant representing the population size, `K`, is set to 2000. A mutation rate (per base position, per generation) is set to $10^{-8}$. The type of mutations to be modelled is configured with a name (`m1`), a given dominance coefficient, fixed (`"f"`) distribution of fitness effects and a given selection coefficient. The type of genomic elements to be simulated is similarly configured with a name (`g1`) and a type of mutations to utilize (`m1`). A single genomic element of length $10^6$ base pairs is set up to use `g1`, representing one chromosome. A recombination rate for the model is set ($10^{-6}$).

The last stage of model setup occurs in the first generation of the model, in which `K` individuals are created and added to a new subpopulation, `p1`, with an implicit initial sex ratio of 0.5 (half male, half female).

Input

In this study, we only varied selection and dominance coefficients of the modelled mutations, set in the `initialize()` callback. However, when the model is applied in more species-specific investigations other parameters would be varied or added, and these can change over time or across subpopulations.

Submodels

The haplodiploid mode of inheritance is set up in the `reproduction()` callback. To produce `K` / 2 males for the new generation *t*, `K` / 2 females are sampled from the previous generation *t* – 1. Each female undergoes recombination, with breakpoints generated by SLiM based upon the recombination rate and chromosome length. Note that the haploid male offspring are "diploid" (individuals are always diploid in SLiM), but their second chromosome is kept empty when they are produced, using the `addRecombinant()` method provided by SLiM:

```
p1.addRecombinant(strand1 = sampledFemale.genome1,
                  strand2 = sampledFemale.genome2,
                  breaks1 = breaks,
                  strand3 = NULL,
                  strand4 = NULL,
                  breaks2 = NULL,
                  sex = "M");
```

Similarly, K / 2 diploid female offspring are produced for generation $t$ by the sexual reproduction of K / 2 males and K / 2 females sampled from the $t-1$ generation. The first chromosome of the diploid females results from the recombination of the mother's genome, and the second chromosome is a copy of the father's haploid genome:

```
p1.addRecombinant(strand1 = sampledFemale.genome1,
                  strand2 = sampledFemale.genome2,
                  breaks1 = breaks,
                  strand3 = sampledMale.genome1,
                  strand4 = NULL,
                  breaks2 = NULL,
                  sex = "F")
```

The sampling of individuals between generations is weighted based on their relative fitness and performed with replacement, so that some adults from generation $t-1$ may produce multiple offspring while others may produce none. The model's `fitness()` callback overrides SLiM's default fitness calculations when modelling haploids. SLiM assumes that all individuals are diploid; in our model, the second genome of haploid males is simply kept empty. When calculating fitness based on the mutations present in an individual, SLiM would thus consider all males to be heterozygous for their mutations. To account for this, the `fitness()` callback assigns males a fitness effect $W = 1 + h_d \times s$, where $s$ is the selection coefficient for the mutation being simulated and $h_d$ the dosage compensation coefficient that determines the effect of selection in haploid males. We set $h_d$ to be 1, thus considering selection to act on haploid males in the same was as on homozygous diploid carriers, but this can be changed.

```
fitness(m1) {
	// Females use the standard fitness calculation
	if (individual.sex == "F")
		return relFitness;

	// males (i.e. haploids that have one copy of the mutation)
	// get a different fitness value, using a haploid dosage coefficient;
	// here we use 1.0 as the haploid dosage coefficient, as if haploid males
	// were homozygous diploid for the mutation, but this can be changed
	return 1.0 + 1.0 * mut.selectionCoeff;
}
```

Following the `reproduction()` callback, the individuals from generation $t-1$ have their fitness set to 0 to ensure that they will die. All individuals from generation $t$ have their fitness set to an arbitrarily high number (to avoid SLiM's default mortality mechanism for non-Wright–Fisher fitness-based models; we instead use fitness during reproduction instead). This ensures that all K individuals survive until the next generation $t+1$ regardless of how many deleterious mutations accumulate.

```
early() {
	inds = sim.subpopulations.individuals;
	inds[inds.age > 0].fitnessScaling = 0.0;
	inds[inds.age == 0].fitnessScaling = 1000.0;
}
```

After this, a `late()` callback identifies and removes mutations that have become fixed in the population, as they no longer contribute to relative fitness.

```
late() {
	// find and remove fixed mutations manually, after mortality; necessary
	// because mutations never fix from SLiM's perspective due to haploid males
	subpops = sim.subpopulations;
	genomeCount = sum(subpops.individualCount) + sum(subpops.firstMaleIndex);
	mutCounts = sim.mutationCounts(NULL);
	fixedMuts = sim.mutations[mutCounts == genomeCount];
	if (fixedMuts.size() > 0) {
		// this code emits a warning about fitness values; it is OK, because
		// fitness in this model is relative fitness, and removal of fixed
		// mutations does not change relative fitness anyway
		suppressWarnings(T);
		subpops.individuals.genomes.removeMutations(fixedMuts, T);
		suppressWarnings(F);
	}
}
```

Note that by summing `subpops.individualCount`, which represents the total number of individuals, and `subpops.firstMaleIndex`, the total number of females, gives us the total number of genomes in the population. The `T` argument of `removeMutations()` creates `Substitutions` objects that are later accessed by `sim.substitutions.size()` to count the number of fixed mutations every 100 generations.

### Supplementary References


Grimm V, Berger U, Bastiansen F, Eliassen S, Ginot V, Giske J, Goss-Custard J, Grand T, Heinz SK, Huse G et al. (2006). A standard protocol for describing individual-based and agent-based models. *Ecol Model.*, 198:115–126.
